# Comparing algorithms for assessing upper limb use with inertial measurement units

**DOI:** 10.1101/2022.02.24.481756

**Authors:** Tanya Subash, Ann David, StephenSukumaran ReetaJanetSurekha, Sankaralingam Gayathri, Selvaraj Samuelkamaleshkumar, Henry Prakash Magimairaj, Nebojsa Malesevic, Christian Antfolk, SKM Varadhan, Alejandro Melendez-Calderon, Sivakumar Balasubramanian

## Abstract

The various existing measures to quantify upper limb use from wrist-worn inertial measurement units (IMU) can be grouped into three categories: (a) Thresholded activity counting, (b) Gross movement score and (c) machine learning. While machine learning algorithms are a promising approach to detect upper limb use, there is currently no knowledge of the information used by these methods, and the data-related factors that influence their performance. A comparison of existing methods was carried out using data from a previous study which was collected from 10 unimpaired and 5 hemiparetic subjects, with annotation to identify periods of functional and non-functional upper limb use. Intra-subject random forest machine learning measures were found to classify upper limb use more accurately than other measures. The random forest measure uses information about the orientation and the amount of movement of the forearm to detect upper limb use. The types of movements and the proportion of functional data included in training/testing set influences the performance of machine learning measures. This study presents the first step towards understanding and optimizing machine learning methods for upper limb use assessment using wearable sensors.

## Introduction

Accurate evaluation of the real-world impact of a neurorehabilitation intervention is crucial to gauge its true value. Thus, there is a growing interest in quantifying how much and how well patients use their affected upper limb(s) outside of therapy. The shortcomings of current questionnaire-based assessments of upper limb use in daily life [1] have led to a surge in the use of wearable sensors for this purpose. Several research groups have explored different sensing modalities [2]–[9] and data analysis techniques [6], [8], [10]–[13] for assessing the amount and quality of upper limb use outside the clinic.

Among the various constructs associated with upper limb functioning in daily life, the most fundamental one is the *upper limb use* – a binary construct indicating the presence or absence of a voluntary, meaningful movement or posture [13]; it is required for deriving other constructs in upper limb functioning [13]. Upper limb use assessment focuses only on measuring willed movements or postures of functional significance. Identifying such movements is a relatively trivial task for a human observing a subject performing various movements. A human’s ability to relate to the movements being observed allows him/her to make judgements about the nature of a subject’s movements. However, doing this in an autonomous manner using technology can be challenging.

The information gathered from measurement systems during everyday life from community-dwelling patients is limited due to constraints on the sensors’ size, wearability, ergonomics, and cosmetics. The most popular sensing modality is inertial measurement units (IMUs) in the form of a wristband [6], [7], [14], which measure linear acceleration and angular velocities of the forearm. To quantify upper limb use from wrist-worn IMU data, various measures have been developed [6], [8], [10]–[12]. These can be grouped into three types: (a) Thresholded activity counting [6], [10], [11], (b) Gross movement score [12], and (c) machine learning [8]. Currently, the threshold activity counting measures are the most popular approach used for quantifying upper-limb use [6], [10], [11]. These measures use quantized linear acceleration (often gravity-subtracted) to compute ‘activity counts’, which is then thresholded to quantify the presence or absence of a functional movement at any given time instant. The gross movement (GM) measure proposed by Leuenberger et al. [12] uses estimates of forearm orientation to decide on the functional nature of upper limb movements. Bochniewicz et al. [15] and Lum et al. [8] used accelerometer data and tested different machine learning methods as measure of upper limb use.

A comparison of the performance of these different upper limb use measures has also recently appeared in the literature [8], [16]. Lum et al. compared the performance of thresholded activity counting to the different machine learning measures on data collected from 10 unimpaired and 10 stroke survivors. They found that the random forest classification algorithm had an overall accuracy of greater than 90%, compared to about 72% for the activity counting methods. They observed that activity counting overestimated upper limb use by indiscriminately picking up both functional and non-functional movements. In our recent study, we made a similar observation comparing the activity counting with the GM measure [16] using data from 10 unimpaired subjects [7]. Activity counting had good sensitivity but poor specificity, compared to GM, which had a lower sensitivity but good specificity for functional movements. A direct comparison of all the existing measures (thresholded counts, GM, and machine learning) of upper limb use is currently missing in the literature. Such a comparison on the same dataset can help delineate the pros and cons of these different measures. This comparison will also help to verify the claims made by Lum et al. [8], and thus, evaluate the generalizability of their results.

Machine learning methods are data-driven approaches. The nature of the training data will impact the performance and generalizability of their result. However, this influence of training data on upper limb use detection has not been investigated in the current literature. Factors such as the relative amount of functional versus non-functional movements and the types of tasks in a dataset can significantly impact a model’s training and testing accuracy. Furthermore, there has also been little work on the interpretability of machine learning methods used for upper limb use assessment. The current methods are black boxes that use a set of handcrafted features to measure upper limb use. Understanding the relative importance of the features could improve the interpretability of these methods. To this end, we also investigated the effect of data-related factors and the importance of different features on the performance of machine learning measures to detect upper limb use.

This study uses an annotated dataset collected from our previous study [17]. This dataset consists of data from 10 unimpaired and 5 stroke survivors performing a set of activities involving arm and hand movements. The current study makes the following contributions to the field,

- Provides a direct comparison of the existing measures of assessing upper limb use and verifies the results presented by Lum et al. [8].
- Compares the performance of common machine learning methods and a classical neural network method.
- Analyses the importance of the features used by current machine learning methods and shows that the essential features have a simple physical interpretation.
- Demonstrates the influence of different data-related factors on the performance of machine learning methods.

## II. Methods

### A. Data Collection

#### 1) Device

Data from a previous study [17] (approved by the institutional review board of Christian Medical College (CMC) Vellore, IRB Min. No. 12321 dated 30.10.2019) on the in-clinic validation of a wrist-worn sensor band, dubbed ‘IMU-Watch’, was used for this analysis. The IMU-Watches log triaxial accelerometer, gyroscope, and magnetometer data at 50 Hz in synchrony.

#### 2) Participants

As part of the study [17], 10 unimpaired individuals and 5 stroke survivors with hemiparesis were recruited. The inclusion criteria for the patients with hemiparesis were: (i) no severe cognitive deficits (Mini-Mental State Examination score (MMSE) higher than 25); (ii) Manual Muscle Test (MMT) grade higher than 2; (iii) age between 25 - 70 years; can actively achieve (iv) at least 30° elevation of the arm against gravity in the shoulder joint with the elbow extended, (v) 20° wrist extension against gravity, and (vi) 10° finger extension (proximal metacarpophalangeal and interphalangeal) of at least one finger against gravity; (vii) ability to open the hand in any position to accommodate a small ball (diameter of 1.8cm) in the palm; and (viii) willingness to give informed consent. Patients were recruited through the inpatient Occupational Therapy unit of CMC Vellore.

The inclusion criteria for unimpaired controls were: (i) no prior history of upper limb movement problems due to neurological conditions; (ii) no current difficulty in upper-limb movements; (iii) age between 25 and 70 years; and (iv) willingness to give informed consent. Subjects who had pain while moving the upper limb and/or allergy to the plastic material used for the IMU-watch casing and straps were excluded from the study.

#### 3) Tasks

Participants performed functional tasks (listed in Table 1) while wearing an IMU-Watch on each arm. The recordings for the tabletop and non-tabletop tasks were taken in two separate sessions, each lasting no more than 15 minutes. Control subjects performed all the tasks, but stroke survivors only completed a subset of these tasks because of difficulty in performing some tasks. Subjects mimicked movements 3-4 times for tasks such as eating or drinking.

**Table 1.** Tasks Performed While Wearing The Imu Watches.

| Tabletop Tasks |  |
| --- | --- |
| Write using a pen | Type on a keyboard |
| Make a call using a mobile phone | Button a shirt |
| Drink from a glass | Drink from a teacup with handles |
| Open a bottle | Comb your hair |
| Wipe a table | Fold a towel |
| Eat from a plate using your hands | Eat from a bowl using a spoon |
| Non-Tabletop Tasks |  |
| Walk 25 meters | Open a door |
| Hit a light switch |  |

#### 4) Ground truth Labelling

The entire experiment was videotaped using a webcam connected to a PC time-synchronized with the IMU-Watches. Two therapists annotated the videos twice with one week gap between each annotation. The annotators were instructed to comply with FAABOS [18], which classifies movements into four classes, based on the functional nature and task-relatedness of the movements, namely, task-related functional, non-task-related functional, non-functional, and no activity or movement. For this analysis, the annotators were asked to reduce the four classes to a binary classification indicating functional and non-functional movements. In addition to the FAABOS annotations, the data was also marked to identify periods of predominantly ‘Hand’ movements, predominantly ‘Arm’ movements, and ‘Non-Functional’ movements to reflect the type of functional movements carried out by the participants during the experiment. The epochs corresponding to the different tasks were also annotated in the data.

#### 5) Dataset Preparation

The final dataset used for the analysis was prepared in the form of a table with the columns and rows corresponding to different features and time stamps, respectively. The different columns of this dataset include the following for each arm.

##### Re-sampled sensor data

Triaxial accelerometer, gyroscope, and magnetometer was recorded approximately at 50 Hz. The raw data from the watches were re-sampled to 50 Hz using zero-order hold interpolation to account for any missing data.

##### Yaw and pitch angles of the IMUs

Yaw and pitch angles of the forearm are estimated from the raw 50 Hz data with respect to an earth-fixed reference frame using the Madgwick algorithm [19]. Offset correction was done on the raw gyroscope data before using the Madgwick algorithm by first identifying “rest” and “move” periods. A rest period is at least 10s long where the signal variance is less than 0.15 deg/s on each gyroscope axis. The mean angular velocity in each axis during a rest period was computed as a gyroscope offset value. This offset value is subtracted from the raw gyroscope data, starting from the current rest period until the next rest period to reduce gyroscopic drift. A fifth-order median filter was applied to the accelerometer data to remove sharp jumps and outliers.

##### Annotations

Four columns that correspond to functional use annotation marked twice by two annotators, were included. Two other columns indicate the tasks and arm/hand use. All annotations were saved at the video frequency of 30 Hz. The annotations were included in the dataset after up-sampling them to 50 Hz using zero-order hold interpolation.

Each row in the dataset is time-stamped. The dataset also includes a column with subject identifiers to differentiate data between different subjects.

### B. Comparison of Measures

The current study compared different measures (shown in Table 2) reported in the literature, along with two new measures (GMAC and MLP). The data processing pipeline for these algorithms was implemented with an interest to remain true to the original work. Where the source material lacked sufficient detail, the processing steps and their associated parameters were chosen empirically as detailed below. The outputs of the different measures were compared with the manual annotations which served as ground truth.

**Table II.** Upper Limb Use Measures That Were Compared

| Measure | Proposed by |
| --- | --- |
| Thresholded Activity Counting | Bailey et al. (Vector Magnitude) [6], De Lucena et al. (Activity Counts) [11] |
| Gross Movement | Leuenberger et al. [12] |
| Hybrid GM and TAC | In the current study |
| Machine Learning | Lum et al. (Random Forests) [8], Support Vector Machine (SVM) [8], Multi-Layer Perceptron (MLP) |

The performance of the different measures across different subjects (and different iterations for intra-subject models) are presented as the ‘sensitivity’ vs. ‘1-specificity’ plots. The measures were also compared using the Youden index [20], a measure of the distance between the top-left corner and the position of the model in the plot, which is given by,

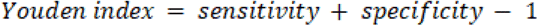

The Youden index for the ideal classifier is 1 and is 0 for a random classifier.

#### 1) Thresholded Activity Counting (TAC)

The amount of acceleration is thresholded using a measure-specific threshold to estimate upper limb use. The computational simplicity of this measure makes it a quick and popular approach [6], [10], [11]. However, while an increased amount of acceleration most likely correlates with increased upper-limb use, the feature is not unique to functional movements, and thus overestimates upper limb use.

##### a) Activity Counts

This measure was proposed by de Lucena et al. [11]. The effects of gravity are removed from the accelerometer data using the 9DOF IMU data with the Mahony algorithm [21]. The magnitude of the gravity-subtracted acceleration data is then bandpass filtered between 0.25 Hz and 2.5 Hz using a 4^th^ order Butterworth filter. The magnitudes are down-sampled to 1 Hz by taking its mean using non-overlapping 1 s bins. The counts are computed by quantizing the magnitudes by 0.017g. A laterality index is calculated using the counts from both arms and thresholded to produce binary signals that indicate use and non-use of both arms. Laterality greater than −0.95 denotes dominant or non-affected arm use, and an index lesser than 0.95 denotes non-dominant or affected arm use.

##### b) Vector Magnitude

This measure proposed by Bailey et al. generates counts at 30 Hz using the Actigraph Activity Monitor [6]. In the current study, the counts were generated from raw acceleration data re-sampled to 30 Hz to match Actigraph sampling rate. This enabled direct application of the methods presented by Brønd et al. [22]. The proprietary actigraph filter was substituted for the Madgwick filter used with 6DOF IMU data. The gravity corrected acceleration data were bandpass filtered between 0.25 Hz and 2.5 Hz using a 4^th^ order Butterworth filter and re-sampled to 10 Hz. The data was then dead-band filtered using the thresholds ±0.068g and summed for every 1s bin. The 2-norm of the acceleration vectors were then computed. A moving average filter with a window size of 5s with a 4s overlap was applied, resulting in the counts at 1 Hz. The counts were filtered using a zero threshold to obtain the binary signal corresponding to upper-limb use.

#### 2) Gross Movement (GM) measure

GM measure [12] is computed for moving windows of 2s with a 75% overlap, resulting in upper limb use estimates at 2 Hz. The yaw and pitch angles were computed using the Madgwick algorithm from the raw acceleration and gyroscope data. If in a 2s window, the overall absolute change in yaw and pitch angles is higher than 30° and the absolute pitch of the forearm is within ±30°, GM is defined as 1, else it is 0. The GM measure exploits the nature of most functional movements to occur in this ‘functional space’.

#### 3) Hybrid GM and TAC (GMAC)

The TAC measures are known to be highly sensitive while having very low specificity, and GM is highly specific but not sensitive [16]. The proposed hybrid measure combines the essential elements of TAC and GM measures as a compromise between them. It combines the counts using the vector magnitude measure with a modified GM; the counts were used instead of the absolute change in yaw and pitch angles. Counts were generated at 1 Hz, and the mean pitch is computed for every 1s bin. Upper limb use is defined as 1 when mean pitch is within ±30° and counts is more than 0, else it is 0.

#### 4) Machine Learning

The TAC and the GM measures are designed to exploit the differences in functional and non-functional movements as measured by an IMU and intuited by a human observer. In currently limited work using machine learning methods for detecting upper-limb use. Lum et al. presented a comparative analysis of supervised machine learning methods and settled on a random forest method [8].

The machine learning methods train models using the ground truth data, i.e., the ‘functional use’ annotations by the human therapists. At each time instant, a single ground truth label was derived as the majority of the four ground truth labels corresponding to two markings by two annotators. In the case of a tie, the instant was labeled as non-functional. We trained and tested two types of models, inter-subject and intra-subject models, with different machine learning methods.

##### a) Features

Eleven features proposed by Lum et al. [8] were computed from the 50 Hz triaxial accelerometer data [8], namely mean and variance for acceleration along each axis, and mean, variance, minimum, maximum, and Shannon entropy of the 2-norm of the accelerometer data. A non-overlapping time window of 0.25s was chosen for computing the features by iterating through different window sizes between 0.25s and 8s and choosing the best-performing window size. The ground truth label for the window was taken from the center, i.e., the label corresponding to time instant 0.125s from the start of the window. A Gaussian kernel with a bandwidth 0.2 was used to compute entropy.

##### b) Machine Learning Methods

A Random Forest (RF), a weighted Support Vector Machine (SVM) with a Radial Basis Function kernel, and a three-layer Multi-Layer Perceptron (MLP) were trained on the contrast, supervised machine learning methods are data-driven and use an annotated dataset to learn the statistical differences between functional and non-functional movements. There is features and ground truth. The model parameters for random forest and SVM (number of estimators for random forest, C and gamma for SVM) were chosen by performing a grid search in a nested cross-validation approach. In nested cross-validation, the hyperparameters were optimized using every fold in the training set as the validation set. The hyperparameters with the best performance were chosen and tested on the testing set.

##### Intra-subject

Stratified 5-fold cross-validation was used to implement the intra-subject model. To account for the variability in performance due to the different random splits, the intra-subject models were generated by iterating through the train-validate-test process 10 times.

##### Inter-subject

Leave-one-out cross-validation was used to implement the inter-subject model.

All classifiers were implemented using the Scikit-Learn package [23].

### C. Interpretation of the Random Forest classifier

To get a handle on the features used by the random forest method, the Gini importance index was computed for the 11 features. The index represents the importance of a feature relative to the other features used in training the model. This analysis was carried out only for the random forest because of the availability of the Gini importance score, and because the random forest performed the best among all methods. To further verify feature importance, three reduced models were trained and validated on the dataset again: (i) mean of (1 feature), (ii) mean of *a*_*x*_, *a*_*y*_, and *a*_*z*_ (3 features), and (iii) mean and variance of *a*_*x*_, *a*_*y*_, and *a*_*z*_ (6 features).

The Spearman correlation of the mean and variance features with the variables used by TAC and GM methods (e.g., Euler angles of the forearm and activity counts) was also computed. This was done to understand the physical significance of the features.

### D. Effect of data-specific factors on machine learning method performance

The nature of the dataset used for developing a machine learning method is a crucial factor in determining performance, arguably as important as choosing a classifier itself. The training dataset used must be a representative sample of the activities and tasks typically performed by a subject in daily life; failing this can drastically affect performance. A training dataset must include all the significant tasks usually expected from a human subject during everyday life; movement patterns drastically different from that in the training dataset are likely to result in poor classifier performance. The current study investigated the four combinations of the presence/absence of selected tasks in training and testing datasets. The tasks chosen for this analysis were opening a bottle, drinking from a cup, and walking (including walking for 25m, hitting a switch, and opening a door); walking tasks were excluded from the patient data since only two patients had performed them. Additionally, the data segments in between tasks marked as unknown tasks were excluded from this analysis.

### E. Statistical analysis

The results from the different analyses were compared using a one-way ANOVA, and t-tests with Bonferroni correction were performed to examine pairwise differences. The following tests were performed:

- Comparison of the Youden indices of different types of measures were done by grouping them into three categories: traditional (AC, VM, GM and GMAC), inter-subject machine learning, and intra-subject machine learning measures.
- Comparisons of the Youden indices of reduced and full models using only the random forest inter- and intra-subject models.
- Comparison of the sensitivities and specificities of all combinations of presence/absence of a task using only the random forest intra-subject models.

The full dataset used in this study and the code for the analysis are available at https://github.com/biorehab/upper-limb-use-assessment.

## III. Results

### A. Which is the best measure for assessing upper limb use?

The performance of the different measures computed across different subjects (and different iterations for intra-subject models) are presented as the ‘sensitivity’ vs. ‘1-specificity’ plots shown in Figure 2A. Figure 2B shows the Youden indices for the different measures. A significant difference was observed between the different types of measures (Fig. 2C) (F = 342.5, p < 0.0001).

**Fig. 1.**
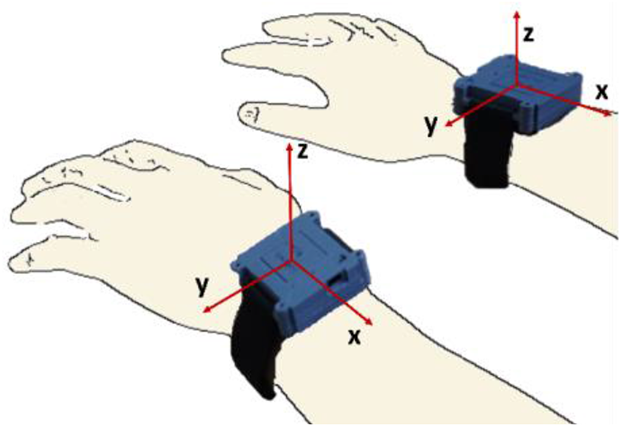
IMU watches with the axes of the accelerometers

**Fig. 2.**
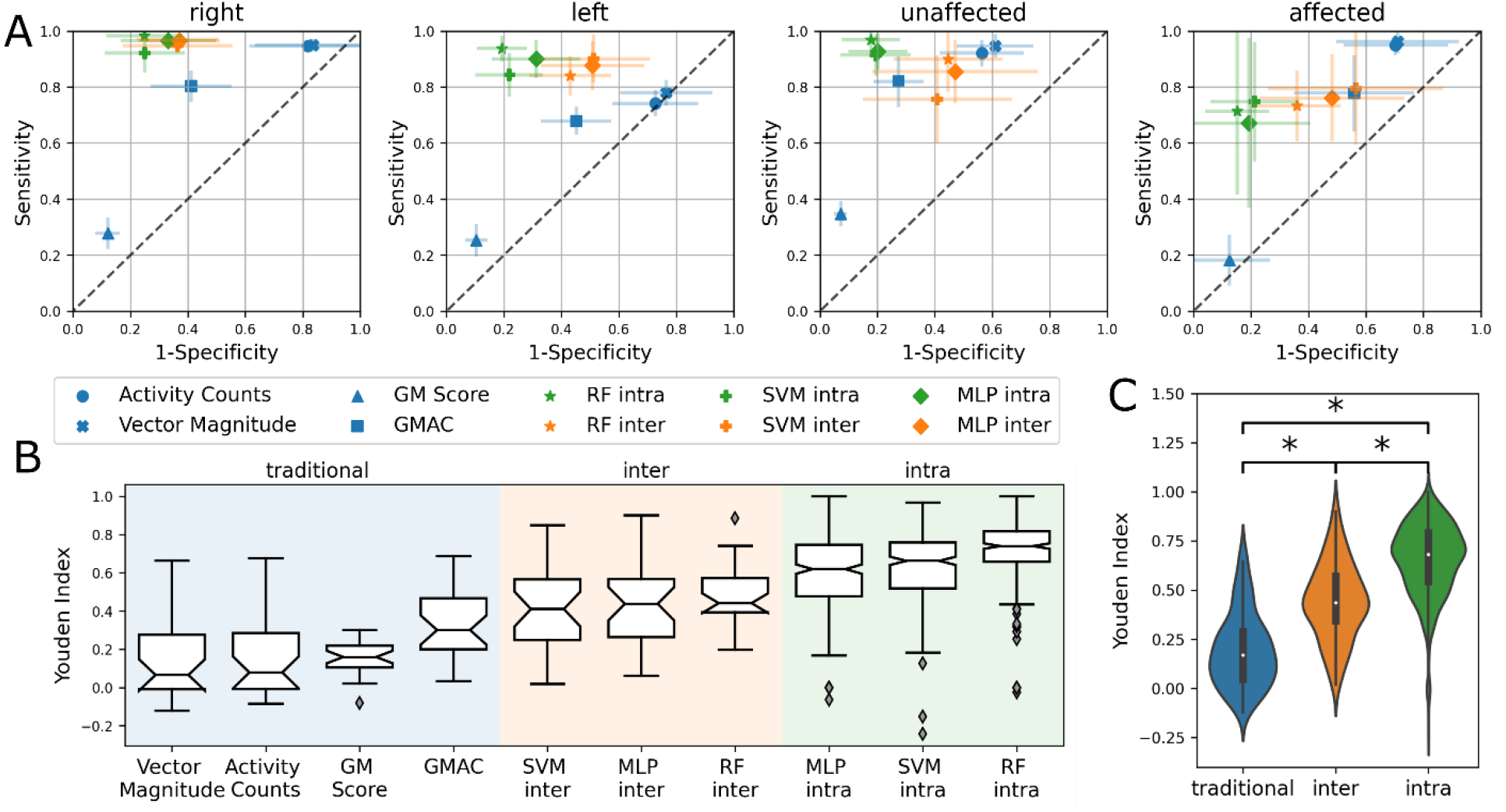
A) Sensitivity vs ‘1-Specificity’ plots depicting the performance of the different measures. The closer a measure is to the top-left corner, the better its performance. The diagonal dashed gray line depicts the performance of a random classifier. B) Boxplot showing the Youden indices for the measures. C) Statistically significant difference between traditional, inter-subject machine learning and intra-subject machine learning measures. *Significant difference (p < 0. 05).

The thresholded activity counting measures had high sensitivity but low specificity with a median Youden index around 0.07. The GM measure had low sensitivity but high specificity and a median Youden index of 0.16. Thus, confirming that thresholded activity counting measures overestimate, while GM underestimates upper limb use. GMAC was found to be a reasonable compromise between thresholded activity counting and GM, resulting in a median Youden index of 0.3.

The machine learning-based upper limb use measures perform better than the traditional measures; intra-subject measures have the best performance (Figure 2C). The random forest intra-subject methods had the highest median Youden index of 0.74 among the machine learning measures. Similar to the findings by Lum et al. [8], the random forest models seem slightly better suited for this application than the SVM or the MLP models. In general, intra-subject models perform better than inter-subject models, which is likely due to inter-subject variability; the inter-subject models for patients are worse than healthy controls (Figure 2A).

### B. What does the random forest classifier do?

The Gini importance index for the mean and variance of *a*_*x*_, *a*_*y*_ and *a*_*z*_ was higher than the other features, with the mean of *a*_*x*_ being the most important feature. The Youden indices for the reduced models are depicted in Figure 4A. The intra-subject models and the inter-subject models for controls using just mean of *a*_*x*_ was the only model worse than the rest (p < 0.05), i.e., using only the mean accelerations achieved similar performance to the full intra-subject model (using all 11 features). However, the acceleration variances were required to achieve similar performance in the inter-subject models for patients; models using just mean of *a*_*x*_ and mean of all accelerations showed statistically significant difference when compared to the full model (p < 0.05). The SVM and MLP methods showed similar trends.

**Fig. 3.**
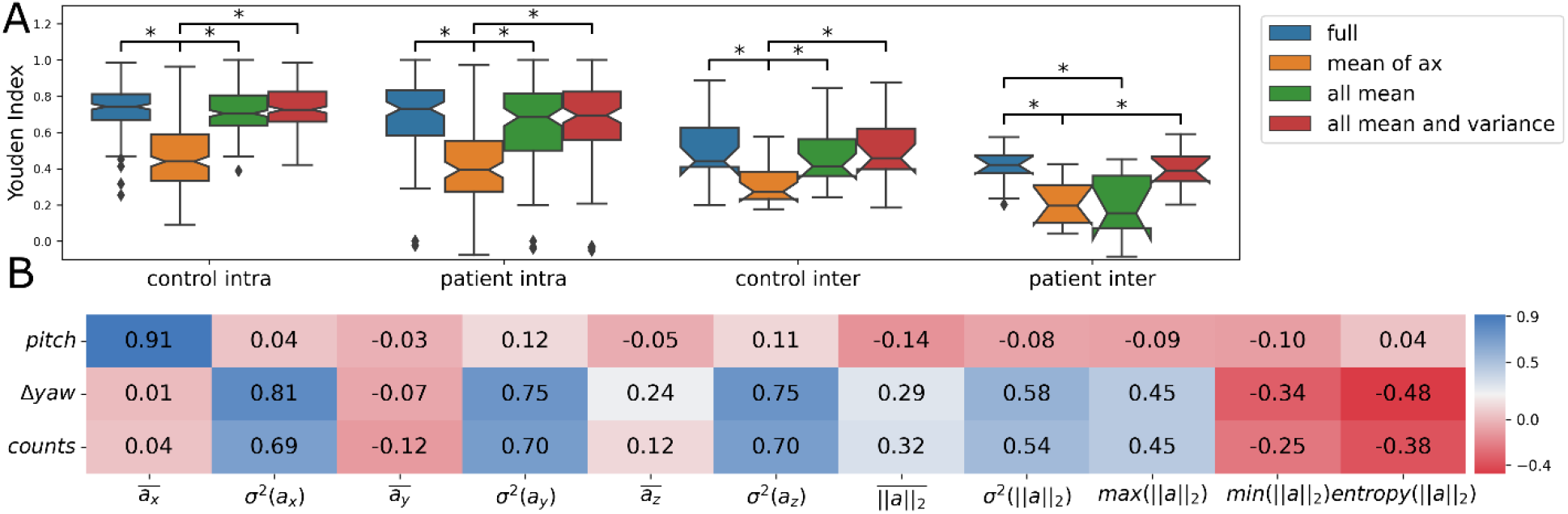
A) Boxplot depicting the Youden indices for the reduced models, *Significant difference (p < 0.05), B) Correlation coefficients between features of the random forest classifier and parameters of GM and TAC

**Fig. 4.**
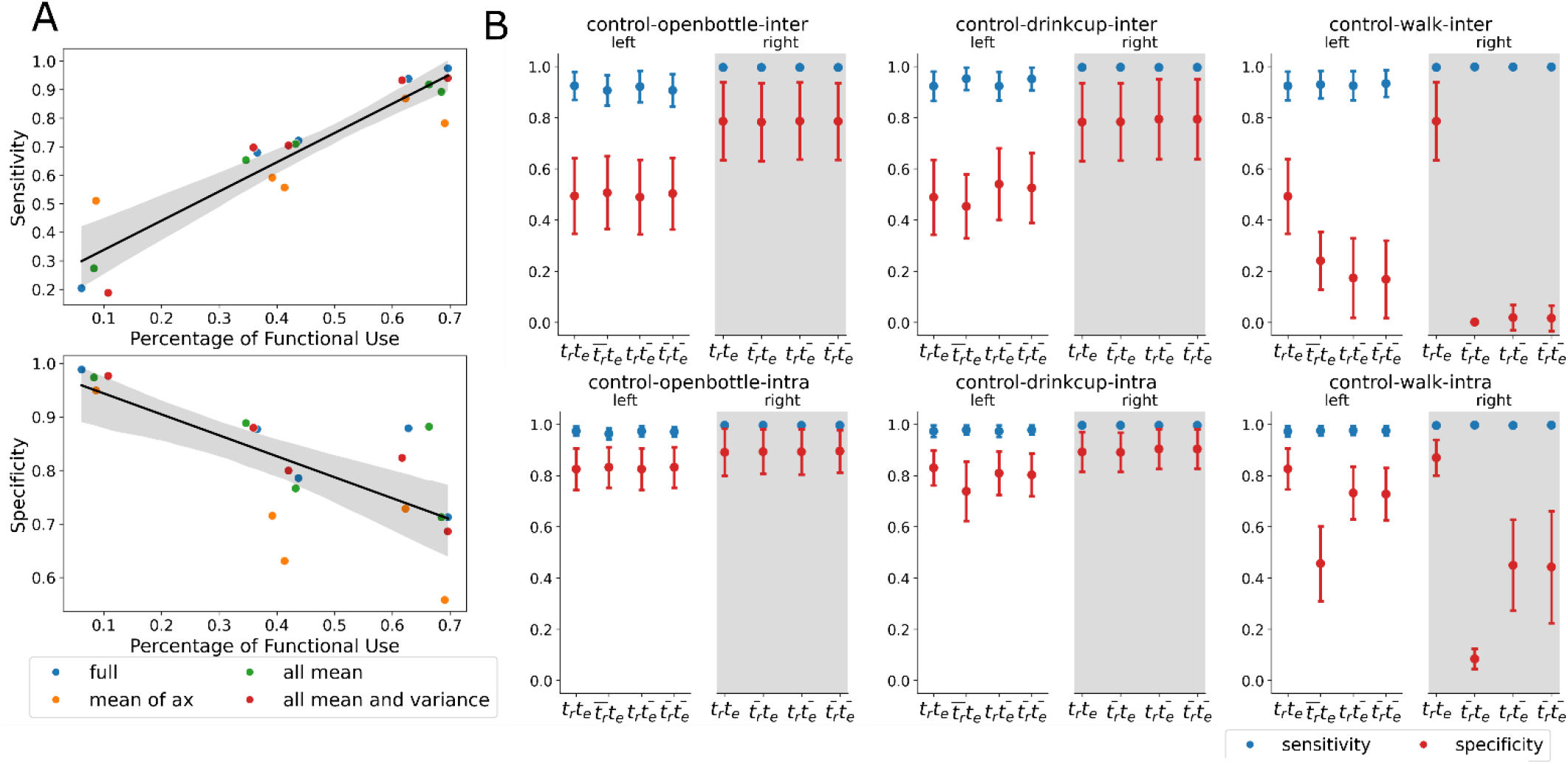
A) Sensitivity and specificity obtained from intra-subject RF classifiers, B) Sensitivity and specificity obtained when certain tasks were removed from the train and test sets. Changes in sensitivity were found to be statistically insignificant in all tasks except in intra-subject models for opening bottle task for the left hand (F = 6.05, p = 0.0005). amount of functional and non-functional movement data can significantly improve performance.

What do the mean and variance features convey? The Spearman correlation coefficient (shown in Figure 4B) between the forearm pitch angle and mean of *a*_*x*_ was 0.91. The coefficient between activity counts and change in forearm yaw angle was around 0.7 with the variance of *a*_*x*_, *a*_*y*_, and *a*_*z*_. Forearm pitch angle indicates forearm’s orientation with respect to ground, while counts and change in yaw indicate the amount of movement in a time window. Therefore, it can be concluded that the random forest method uses a mix of information used by the traditional TAC and GM measures to detect upper limb use. The high performance achieved by only using mean *a*_*x*_ suggests the forearm pitch plays a significant role in determining upper limb use, at least in the current dataset.

### C. Does a gyroscope improve classifier performance?

Features from the IMU’s gyroscope (e.g., mean and variance of *g*_*x*_, *g*_*y*_ and *g*_*z*_) were computed like the other accelerometer features. However, adding gyroscope features did not show statistically significant improvements (p > 0.1) in detection performance.

### D. How does the nature of the dataset affect a machine learning method’s performance?

There are two data-related aspects that can impact performance. The first is the effect of the proportion of the two classes of movements (functional versus non-functional), and the second is the effect of the tasks present in the dataset.

#### 1) Proportion of functional and non-functional data

The performance variance of the machine learning methods was high for the affected arm of patients due to differences in the amount of functional use in the data for the different patients. The sensitivity and specificity of the intra-subject machine learning models for a patient were proportional to the amount of functional and non-functional movements in the dataset, respectively. This is depicted in Figure 4A for the patient data. Thus, training on a large dataset with an equal amount of functional and non-functional movement data can significantly improve performance.

#### 2) Presence/absence of tasks in testing/training datasets

Figure 5B shows the sensitivity and specificity of the random forest method (with all 11 features) employing different train and test datasets. The presence of a particular task in the train set is denoted by t_r_ and its presence in the test set is denoted by t_e_; their absence is denoted by 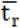 and 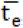, respectively. The following observations were made from the results obtained (Figure 5B).

i. There aren’t significant changes in the sensitivity under the different conditions for any of the tasks.
ii. Specificities for the functional tasks opening a bottle and drinking from a cup do not change much under the different conditions. However, there is a drop in specificity under 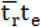 for drinking from a cup for the intra-subject model for the left-hand data (F = 19.1, p < 0.0001). This could be because almost all control participants used only their right hand for this task, resulting in mostly non-functional data in the left-hand data.
iii. There is a significant drop in the specificity for walking tasks under 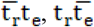 and 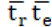 particularly for 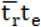 when compared to the baseline (t_r_t_e_)(p < 0.0001).
iv. The results obtained for the patient data at the individual level were varied. Each patient performed different and fewer tasks, and so, meaningful conclusions about the tasks in the dataset could not be made.

## IV. Discussion

The current study compared the sensitivity and specificity of existing measures for quantifying upper limb use – a fundamental construct in assessing upper limb functioning [13]. The results from the comparative analysis show that the machine learning-based measures are better than the other measures presented here and are a promising approach for upper limb use assessment using IMUs. Our analysis, using an independent dataset, confirms the results reported by Lum et al. [8]: (a) the random forest method is slightly better than SVM in detecting upper limb use; we also found that it is also slightly better than an MLP neural network. (b) intra-subject machine learning models are better than inter-subject models, which is due to the inter-subject differences in the movement patterns. Another possible reason for the reduced performance of the inter-subject models could be the small number of subjects included for training. However, it is unclear why random forests perform slightly better than the other machine learning methods. Similar observations about random forests are seen in other applications [24].

Although the improved performance in upper limb use assessment through machine learning methods is valuable in practice, the black-box nature of these methods obscures their mechanism of operation. Traditional measures (TAC and GM), despite their poor performance, offer an intuitive explanation of their classification since they employ interpretable quantities such as counts, forearm pitch, and yaw. Previous work by Lum et al. employed models using 11 accelerometer features and did not attempt to understand the roles of these different features and their physical significance. Understanding the most relevant and important movement features can guide the optimal selection of sensors for measurements and further improve measure performance. The current study shows that the random forest method employs a combination of arm orientation (mean acceleration features) and the amount of movement (variance of acceleration) to estimate upper limb use. Either GM or TAC does not measure up to the random forest method as they do not use all the relevant information and have relatively simple decision rules. However, an advantage of these traditional methods is that they do not require any training, unlike the machine learning methods. The hybrid GMAC measure performed very closely to the inter-subject machine learning models, indicating that it might still be useful in the absence of training datasets required for employing machine learning methods. However, the detection ability of the intra-subject models is unmatched owing to the differences between subjects, especially in those with hemiparesis.

Most functional movements were performed on top of a table in the current dataset, while walking formed a significant portion of the non-functional movements class. Therefore, it is no surprise that the pitch of the arm (mean acceleration) was an important feature in classifying movements. In another dataset where only tabletop tasks are included, the pitch may not play as significant a role, and activity counts could be the determinant in distinguishing between functional movements and rest.

The observations made in this study about the nature of a dataset are preliminary, largely because of the limited number and variety of tasks included in the experiment. The observed results on the impact of the presence/absence of a particular task on a measure’s performance can be explained by two factors: (a) how well represented the movements of this particular task are in that of the other tasks in the dataset, and (b) how much the proportion of the functional class is affected by the presence or absence of a particular task in the dataset. Tasks that have similar movement patterns and similar proportions of functional movements to other tasks can be removed from the dataset without losing performance. But tasks with unique movement patterns and different proportions of functional movements must be included in the training dataset to ensure good performance for a machine learning method. It can be safely concluded that the choice of tasks for the training dataset has an integral part in determining the performance of machine learning-based upper limb use measures. Machine learning methods should ideally be trained on a sample representative of daily life behavior, consisting of similar tasks and proportions of functional and non-functional movements. The variability across subjects in everyday tasks and the amount of functional activity compels the use of intra-subject models for classifying upper limb movements. However, training and deploying these intra-subject models in practice is cumbersome, primarily because of the arduous process of manual ground truth labeling. With a much larger dataset with more participants, it is possible that inter-subject model performance may improve. In which case, pre-trained, generalized models can be deployed without tuning for individual subjects.

The relatively small dataset size used in this current study is its main shortcoming. The dataset contained 10 unimpaired and 5 stroke survivors with hemiparesis carrying out a set of daily activities; stroke survivors did not perform some of the tasks due to difficulty in performing them. There were also variations in the percentages of functional and non-functional movements within the dataset. Nevertheless, the agreement of the results with that of Lum et al. [8] restores confidence in the results, despite the small dataset used.

The use of machine learning-based measures for quantifying upper limb use seems to be the way forward. Future work in this space must focus on improving the performance of machine learning methods to deploy them in clinical practice efficiently. We propose the following two activities that are worth pursuing:

1. Developing a large, open, annotated dataset with unimpaired and people with impaired movement abilities, performing a wide range of daily activities wearing different types of sensors (at least wrist worn IMUs). This can stimulate work on developing and validating optimal classifiers with high accuracy and reliability. The proposed dataset must have a good sample of the types of movements expected from patients in daily life and a good proportion of functional and non-functional movements of interest.
2. Automatic annotation of functional/non-functional movements from video recordings using an RGBD camera must be explored to eliminate the cumbersome manual labelling process. Recent work has shown that data from multiple IMUs can detect different functional primitives of complex upper limb movements [25]. Thus, it is likely that pose estimates obtained from an RGBD camera can be used for automatically classifying functional and non-functional movements, along with different task types as well.
3. We believe that exploring these two avenues will help make sensor-based upper limb use detection highly accurate and efficient for routine clinical use.

## V. Conclusion

This paper presented a detailed comparison of existing measures to quantify upper limb use, and confirms previous finding that an intra-subject random forest measure outperform others. The current work sheds light, for the first time, on the information used by random forest measure, demonstrating that it uses a combination of arm orientation and amount of movement to detect upper limb use. The work also demonstrates the effect of factors related to data, such as class proportion and types of tasks, on the performance of machine learning measures. We strongly believe that this study is a step towards understanding and optimizing machine learning measures to detect upper limb use.

