## Supplementary material for "Comparing algorithms for assessing upper limb use with inertial measurement units"

Gini importance indices of random forest features across all groups (left, right, affected, unaffected)

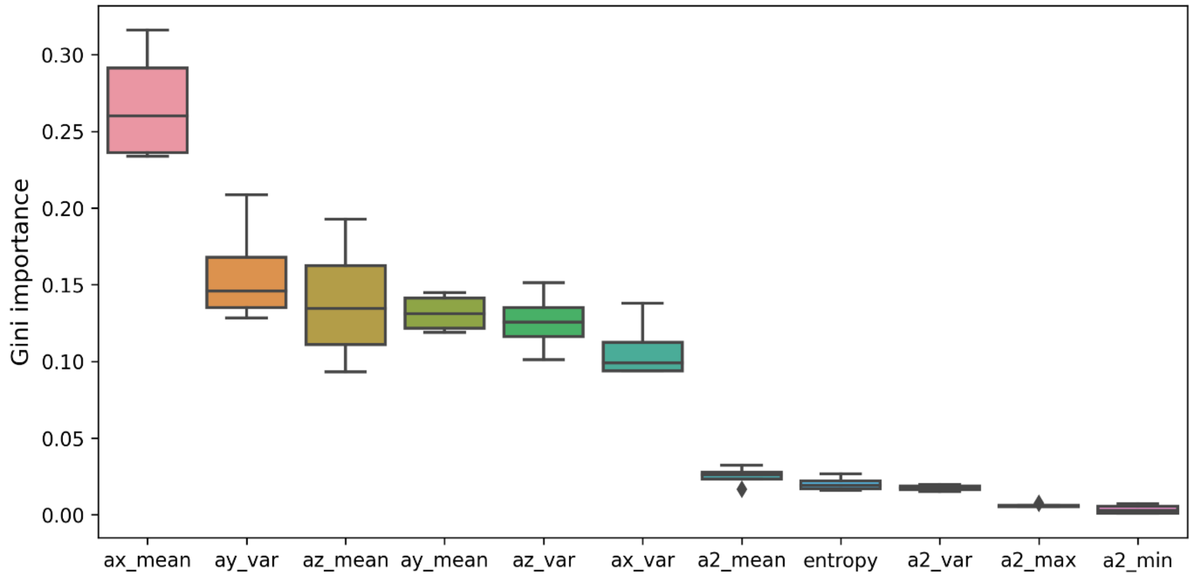


Effect of adding gyroscope features to the random forest

| | | without gyroscope | with gyroscope | F value | P value |
|---|---|---|---|---|---|
| Inter | sensitivity | 0.87 (0.11) | 0.89 (0.14) | 0.38 | 0.54 |
| | specificity | 0.60 (0.16) | 0.64 (0.15) | 1.24 | 0.27 |
| Intra | sensitivity | 0.92 (0.12) | 0.96 (0.03) | 2.44 | 0.12 |
| | specificity | 0.79 (0.08) | 0.77 (0.08) | 0.53 | 0.47 |
